# Queen pheromone modulates the expression of epigenetic modifier genes in the brain of honeybee workers

**DOI:** 10.1101/2020.03.04.977058

**Authors:** Carlos Antônio Mendes Cardoso-Junior, Isobel Ronai, Klaus Hartfelder, Benjamin P. Oldroyd

**Affiliations:** Departamento de Biologia Celular e Bioagentes Patogênicos, Faculdade de Medicina de Ribeirão Preto, Universidade de São Paulo, Ribeirão Preto, Brazil; Behaviour, Ecology and Evolution (BEE) laboratory, Macleay Building A12, University of Sydney, Sydney NSW 2006, Australia

**Keywords:** DNMT, HAT, HDAC, SIRT, honey bee

## Abstract

Pheromones are used by many insects to mediate social interactions. In the highly eusocial honeybee (*Apis mellifera*) queen mandibular pheromone (QMP) is involved in the regulation of reproduction and behaviour of workers. The molecular mechanisms by which QMP acts are largely unknown. Here we investigate how genes responsible for epigenetic modifications to DNA, RNA and histones respond to the presence of QMP. We show that several of these genes are upregulated in the honeybee brain when workers are exposed to QMP. This provides a plausible mechanism by which pheromone signalling may influence gene expression in the brain of honeybee workers. We propose that pheromonal communication systems, such as those used by social insects, evolved to respond to environmental signals by making use of existing epigenomic machineries.

## 1. Introduction

The transition to social living in the eusocial insects required that the reproductive interests of individual workers be subsumed by the collective interests of the colony [1–3]. In particular, workers are functionally sterile, whereas queens are highly fecund [4]. For such systems to evolve it was also necessary that tasks can be distributed among workers in ways that enhance colony-level productivity [5–8]. Both, the regulation of worker fertility and the efficient allocation of tasks among workers required the evolution of effective inter- and intra-caste communication systems that can rapidly respond to the changing needs of the colony. Communication between nestmates most often occurs via pheromones [9,10], chemical signals that are produced by one individual, and cause changes in the behaviour or physiology of another [11,12].

In the western honeybee (*Apis mellifera*) a key pheromone is the queen mandibular pheromone (QMP), which is a blend of fatty acids secreted by the head glands of the queen. It affects several important traits of workers, including their reproduction [13], retinue response to queens [14,15], learning capacity [16], nestmate recognition [17] and age at onset of foraging [18]. Phenotypic variation in these traits is associated with differential gene expression in the brains of workers [19,20]. Nonetheless, little is known about the intermediate steps between QMP production and release by the queen, the regulation of gene expression in workers, and changes in their behaviour [21–24].

Epigenetic mechanisms are a likely mediator between a worker’s social environment and global gene expression responses [25]. Several classes of epigenetic mechanism described in honeybees are potentially associated with environmental cues [26–29]. For example, DNA methylation, a reversible chemical modification of cytosines in CpG contexts, is associated with behavioural maturation in the brains of honeybee nurses and foragers [30,31]. DNA methylation is catalysed and maintained by the DNA methyltransferase (DNMT) family of enzymes [32,33]. Interestingly, in honeybees, the expression of genes associated with the maintenance of DNA methylation levels after DNA replication (*Dnmt1a* and *Dnmt1b*) are modulated by different social stimuli to *Dnmt3,* an enzyme that establishes DNA methylation patterns *de novo* [25,34–36]. In addition, the expression of *Dnmt2* (also called *Trdmt1*), a gene whose enzyme product methylates RNA substrates [37], is affected by different social contexts [25,30]. These studies suggest that epigenetic machineries associated with nucleotide modification are affected by several environmental cues.

Another epigenetic mechanism, histone post-translational modifications (HPTMs), change chromatin structure by altering the physicochemical affinity between DNA and histones and thereby affect gene expression [38]. HPTMs are catalysed by histone modifier proteins [33], which can be divided into three functional classes: writers, erasers and readers. “Writer” enzymes add chemical radicals to histone tails by covalent modification. For example, lysine acetyltransferases (KATs) promote acetylation of lysine residues [33], which reduces the affinity between DNA and nucleosomes. Histone acetylation induces chromatin relaxation and is often associated with increased gene expression [39]. In contrast, “Eraser” enzymes remove such chemical radicals from histone tails. Classical eraser enzymes are the histone deacetylases (HDACs) and Sirtuins, which remove acetyl groups from lysine residues, resulting in chromatin compaction and, consequently, inhibition of gene expression [33,40]. Finally, “Reader” enzymes recognise epigenetic modifications and induce chromatin remodelling through the recruitment of protein complexes [33]. A honey bee proteome study [27] has shown that histone tails are extensively modified by epigenetic marks, indicating that writer, eraser and reader enzymes are present in the honey bee. Furthermore, differential accumulation of HPTMs has been associated with caste differentiation and behaviour in bees and ants [27,41–44].

Given that QMP affects behaviours in honey bee workers [14,15,18,45] we hypothesised that the expression of genes associated with epigenetic modification to nucleotide and histones would respond to QMP exposure in the brain of honey bee workers. These epigenetic mechanisms can, thus, serve as proxies to understand the regulation of global changes in gene expression in a complex social environment.

## 2. Material and Methods

### (a) Biological material

To obtain age-matched adult workers we collected brood frames from four queenright *A. m. ligustica* source colonies and kept them in an incubator overnight at 34.5 °C. From each source colony, workers were randomly allocated to two cages (n = 150 bees per cage, eight cages in total). One cage from each colony (QMP^+^) was furnished with a 0.5 queen equivalent per day QMP strip (Phero Tech Inc. Canada), which is an effective queen mimic in cage experiments with young workers [21,46]. The other cage from each colony (QMP^-^) contained no QMP strip. Pollen, honey and water were provided *ad libitum*. Food was replenished when necessary, and the number of dead workers was recorded each day, which was nearly the same in the QMP^+^ and QMP^-^ cages (data not shown). Cages were kept in an incubator at 34.5 °C for four days. Workers were collected on dry ice at Day 0 (directly from the brood comb), Day 1 and Day 4. Day 1 was chosen to identify genes with a quick response to the QMP treatment, and Day 4 was chosen to identify the genes that are still influenced by the QMP exposure after prolonged exposure. We then dissected the brains of the workers on dry ice [47].

### (b) Identification of the honeybee DNA methyltransferases and histone modifiers

We identified the nucleotide and histone modifier genes in the honey bee genome (Amel_4.5) [48], searching manually for the names of each epigenetic gene in GenBank (NCBI Resources Coordinators 2018) (Table S1) based on a large list of histone-modifier genes present in eukaryotes [50,51]. We filtered this list by selecting those associated with acetylation and deacetylation processes. From this list we identified the proteins that are predicted to reside in the nucleus using ProtComp v9.0 (Softberry, Inc.). The genes and their respective proteins were characterised following a previously described workflow [13].

### (c) Gene expression quantification and bioinformatics analysis

Each sample consisted of a single brain. We extracted total RNA from the brain through maceration in TRIzol (Invitrogen) and a Direct-zol™ RNA Miniprep kit (Zymo Research). The RNA was treated with Turbo DNase (Thermo Fisher Scientific) and quantified with a Qubit 2.0 Fluorometer (Invitrogen). cDNA was synthesised from 600 ng of RNA using a SuperScript™ III Reverse Transcriptase Kit (Invitrogen) with oligo(dT) primer and suspended in ultrapure water (5 ng cDNA/μL).

The expression of four nucleotide modifier genes and 11 histone modifier genes (Table S1) was quantified by reverse transcription quantitative real-time PCR (RT-qPCR) [46,52]. Assays were set up with 2.5 μL SsoAdvanced(tm) Universal SYBR^®^ Green Supermix (Bio Rad), 1.25 pmol of each primer, 1 μL diluted cDNA (5 ng) in a total volume of 5 μL using a CFX384 Real-Time System (Bio-Rad). For each experimental sample (four source colonies, three ages and two treatments) three technical replicates were conducted. Cycle conditions were as follows: 95 °C for 10 min followed by 40 cycles of 95 °C for 10 s, 60 °C for 10 s and 72 °C for 15 s. At the end of the RT-qPCR protocol a melting curve analysis was run to confirm a single amplification peak. Primer efficiencies (Table S2) were calculated based on an amplification curve of 10 points obtained through serial dilution of mixed cDNA samples. The expression of the genes of interest was normalised against the expression of two reference genes (*Rpl32* and *Ef1α*), whose expression was found stable according to *BestKeeper* [53]. Relative expression levels were calculated [52], using a formula that normalises gene expression to the reference genes taking into account the efficiency of each primer set. The genes, primer sequences and efficiencies are listed in Table S2.

### (d) Statistical analysis

To compare the expression of the QMP^+^ and QMP^-^ treatments at Day 1 and Day 4 we used a generalised linear mixed model (GLMM) with ‘colony’ as random effect and ‘treatment’ and ‘age’ as fixed effects. To model the gene expression data, we used link = identity, family = Gaussian. Where necessary, a transformation log_10_ function was applied (see Table S3 for details). We used Day-0 data as a baseline for gene expression. GLMM analyses were performed in R [54] loading the packages lme4, car and emmeans. An adjusted *p*-value (Tukey correction for each gene) lower than 0.05 was considered significant for all statistical tests.

## 3. Results

Using the protein sequences of the 15 genes studied, we first acquired *in-silico* evidence (e.g. subcellular location, predicted domains and homology with other species) that each gene was a *bona fide* epigenetic modifier of DNA, RNA or histones (Table S1). In 1-day old workers the expression of twelve genes associated with epigenetic processes (*Dnmt1b, Dnmt2, Dnmt3, Kat2a, Kat3b, Kat6b, Kat8, Hdac1, Hdac3, Sirt 1, Sirt7* and *Rcs1*) was affected by exposure to QMP (GLMM, *p* < 0.05, Figures 1 and 2, Table S3). However, only four genes (*Dnmt1b, Dnmt2, Kat3b* and *Sirt7*) continued to be differentially expressed at the age of four days (GLMM, *p* < 0.05, Figures 1 and 2, Table S3). Age was statistically significant for 13 of the 15 genes (GLMM, *p* < 0.05, Figure 1 and Table S3), the exceptions being *Kat 7* and *Dnmt3*. A significant interaction between treatment and age was found for three genes: *Hdac1, Sirt1* and *Kat6b* (*p* < 0.05, Table S3).

**Figure 1.**
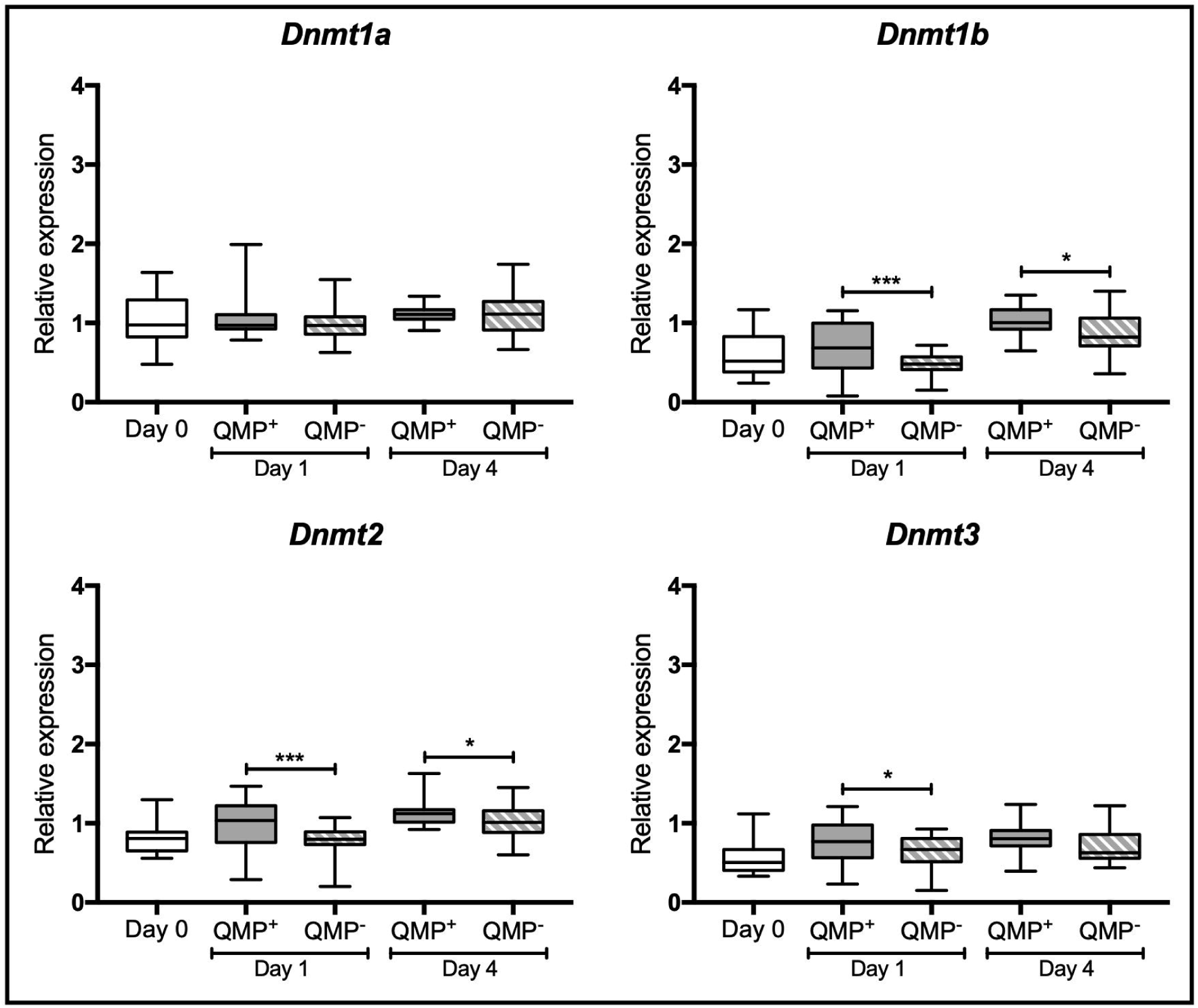
Relative expression of four nucleotide modifier (DNA methyltransferase) genes in the brains of 0-, 1- and 4-days old honeybee workers, exposed to queen mandibular pheromone (QMP^+^) or not (QMP^-^). Each box shows the interquartile range (25^th^-75^th^ percentiles) and the median (line), while whiskers represent the farthest points of 2.5^th^-97.5^th^ percentiles. Relative expression was calculated for each gene at all three ages. Day 0 was used as the baseline for gene expression. Statistical information: GLMM test with Tukey correction for multiple pairwise comparisons, * *p* < 0.05, ** *p* < 0.01, *** *p* < 0.001, N=32; N=8 from each of the four colonies).

**Figure 2.**
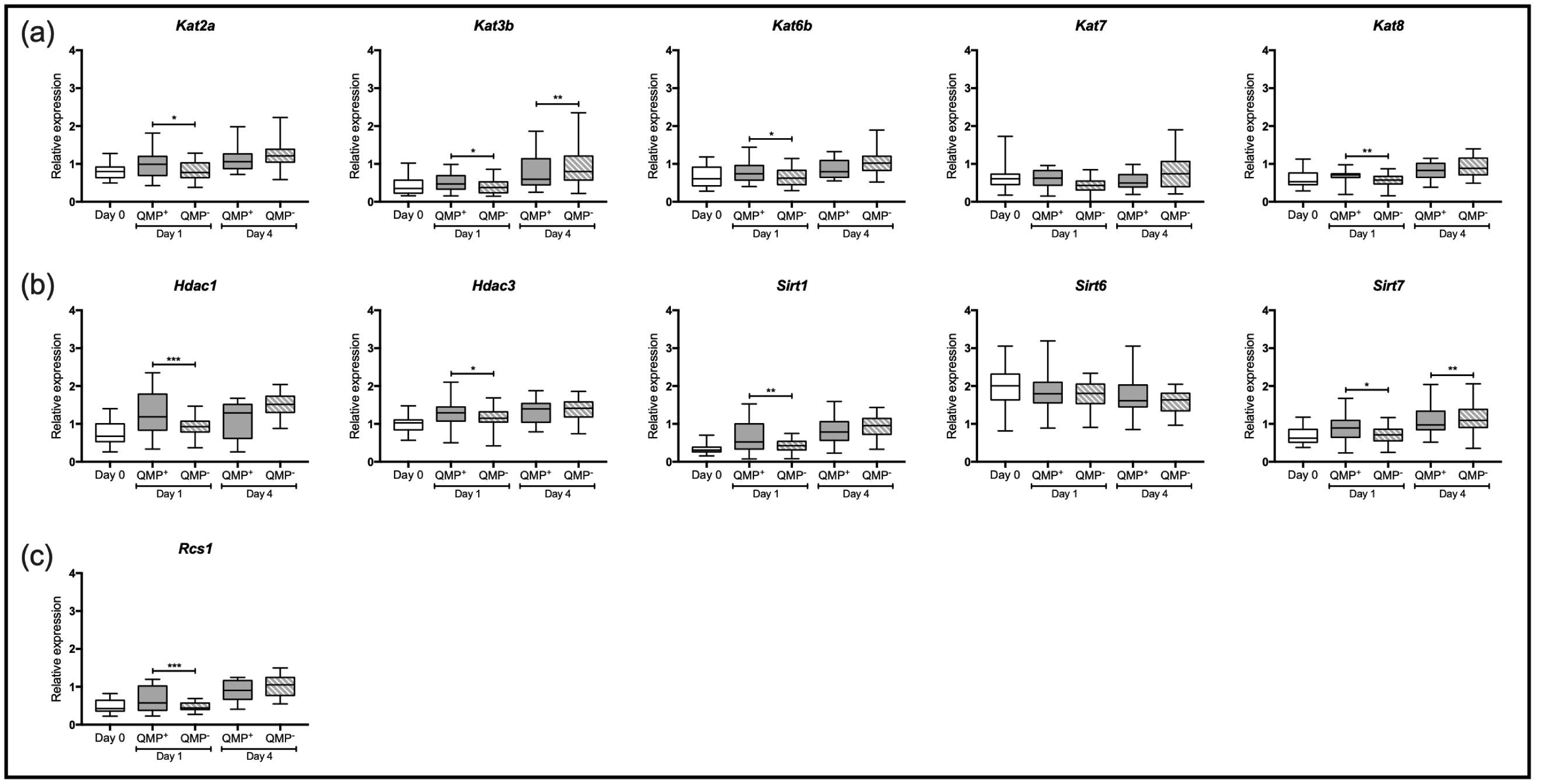
Relative expression of 11 histone modifier genes in the brains of 0-, 1- and 4-days old honey bee workers, exposed to queen mandibular pheromone (QMP^+^) or not (QMP^-^). (a) Relative expression of histone acetyltransferases genes (writer enzymes). (b) Relative expression of histone deacetylases and Sirtuin genes (eraser enzymes). (c) Relative expression of the *Rsc1* gene (reader enzyme). Each box shows the interquartile range (25^th^-75^th^ percentiles) and the median (line), while whiskers represent the farthest points of 2.5^th^-97.5^th^ percentiles. Relative expression was calculated for each gene at all three ages. Day 0 was used as the baseline for gene expression. Statistical information is as in figure 1.

## 4. Discussion

Our study shows that QMP affects the expression of 12 of 15 genes that are associated with epigenetic processes in the brain of honeybee workers. As predicted, our data indicates that epigenetic mechanisms are likely mediators between queen pheromone signalling and the regulation of worker gene expression. Given that QMP alters worker behaviour [14,15,18,45], we suggest that pheromonal communication evolved by making use of existing epigenetic mechanisms that orchestrate transcriptomic changes necessary to propagate pheromonal information.

Some expression responses are particularly worthy of note. For instance, the expression of *Dnmt3,* the *de novo* methylator of DNA, is regulated by queen pheromones in brains of honeybee workers (this study) and whole-body RNA extracts of honeybee workers [35] and two ant species (*Lasius flavus* and *Lasius niger*) [55]. Expression of *Kat 8* is upregulated in the brains of QMP-treated honeybee workers. This gene is differentially spliced in *L. flavus* ants treated with queen pheromone [55]. Together, these results suggest an evolutionary conservation in the epigenetic pathways responsive to queen pheromones in social insects.

We detected that several histone modifiers genes associated with acetylation/deacetylation processes are differentially expressed in the brains of adult workers. This finding suggests that queen signals influence the modification of histones to promote chromatin reorganisation and thereby altering gene expression in worker brains. In line with this hypothesis, histone acetylation contributes to the regulation of foraging behaviour in ants [43]. Interestingly, it was recently shown that honeybee queens regulate worker fertility through polycomb repressive complex 2 (PRC2) activity and differential histone methylation marks [56]. We propose that queens, via QMP, influence modifications to histones to regulate behavioural plasticity in the brains of honeybee workers, just as they do in ovaries.

Pheromonal modulation of gene expression in honeybee workers changes over time [19,20]. Gene expression in QMP^-^ workers is relatively stable from Day 0 to Day 1 when compared to QMP^+^ workers, suggesting QMP *actively* promotes expression of several epigenetic modifier genes already within 24 hours. Only four of these continued to be differentially expressed after 4 days of QMP exposure, indicating that the expression of the majority epigenetic modifiers is dynamically switched on and off [19].

Our study provides evidence that many genes associated with epigenetic modification are differentially expressed in the brains of honeybee workers in response to the presence of queen pheromone. These changes wrought by the genes studied here likely drive changes in gene expression in the brains of adult workers, providing a plausible mechanism by which a queen can influence both the rate of behavioural maturation and reproductive behaviour of her workers. This property of QMP would explain why it acts both as a short-term ‘releaser’ pheromone that merely indicates queen presence, as well as a long-term ‘primer’ pheromone that regulates behavioural maturation and reproductive behaviour.

## Data accessibility

The data that support this study are available in the supplementary material.

## Authors’ contributions

CAM designed the study, carried out the cage experiments, dissections, performed molecular lab work, analysed data and wrote the manuscript draft. IR carried out the protein characterisation and wrote the manuscript. BPO and KH supervised the work and revised the manuscript. All authors gave final approval for publication.

## Competing interests

The authors declare no conflict of interests.

## Funding

This research was funded by the Australian Research Council DP180101696 and Fundação de Amparo à Pesquisa do Estado de São Paulo (FAPESP - #2016/15881-0 and 2017/09269-3 to CAM). The funders had no role in study design, data collection and analysis, decision to publish, or preparation of the manuscript.

## Notes

### Competing Interest Statement

The authors have declared no competing interest.

